# Dorsal and ventral premotor cortices differentially influence contralateral motor cortex excitability

**DOI:** 10.64898/2026.04.29.721139

**Authors:** Elnaz Allahverdloo, Larissa K. Chiu, Amanda O’Farrell, Nesrine Harroum, Numa Dancause, Jason L. Neva

**Affiliations:** École de kinésiologie et des sciences de l’activité physique, Université de Montréal, Montréal, QC, Canada; Centre de recherche de l’institut universitaire de gériatrie de Montréal, Montréal, QC, Canada; Département de neurosciences, Université de Montréal, Montréal, QC, Canada; Centre Interdisciplinaire de Recherche sur le Cerveau et l’Apprentissage (CIRCA), Université de Montréal, Montréal, QC, Canada; Institut Courtois d’innovation biomédicale (CI2B), Université de Montréal, Montréal, QC, Canada

**Keywords:** dorsal premotor cortex, ventral premotor cortex, primary motor cortex, interhemispheric inhibition, ipsilateral silent period, transcranial magnetic stimulation

## Abstract

Dorsal (PMd) and ventral (PMv) premotor cortices can modulate contralateral primary motor cortex (M1) excitability, but their distinct interhemispheric influence via transcranial magnetic stimulation (TMS) remains unclear. Single-pulse TMS over PMd, PMv and M1 assessed transcallosal inhibition via the ipsilateral silent period (iSP). Dual-site TMS examined short-(10 ms inter-stimulus interval [ISI]), long-(50 ms ISI) and non-callosal-(0 ms ISI) interhemispheric inhibition (IHI). An iSP was elicited from PMd, PMv, and M1, with distinctly evoked iSP parameters. The iSP magnitude was greatest from M1, followed by PMd and then PMv, while iSP duration was greatest for M1 and showed no differences between PMd and PMv. Dual-site TMS revealed that PMd and M1 inhibited contralateral M1 excitability across all ISIs, while PMv showed inhibition at 0-and 50-ms ISIs. PMd and M1 demonstrated greater short-IHI compared to PMv, all demonstrating similar long-IHI, and PMd demonstrating greater non-callosal-IHI than M1. PMv displayed distinct IHI across ISIs, PMd showed differences across most ISIs and M1 demonstrated the fewest differences across ISIs. Longer iSP duration related to greater long-IHI magnitude elicited from PMd and PMv. Our findings demonstrate differential IHI from PMd and PMv on contralateral M1, which may inform neuromodulation strategies in rehabilitation contexts.

---

1. **INTRODUCTION**

Goal-directed movements and motor learning rely on a distributed network of motor-related regions, with the primary motor (M1) and premotor (PMC) cortices playing critical, yet distinct, roles (Rizzolatti and Luppino 2001; Hoshi and Tanji 2007; Van Malderen et al. 2023). While M1 is traditionally viewed as the main driver of movement execution and a principal source of corticospinal output, PMC – encompassing its dorsal (PMd) and ventral (PMv) subregions – contributes to motor planning, preparation, and execution, and has its own corticospinal output (Hoshi and Tanji 2004; Chouinard and Paus 2006; Denyer et al. 2023). Critically, PMC can shape M1 output via cortico-cortical connections within and across hemispheres, as well as through subcortical pathways (Baumer et al. 2006; Côté et al. 2017). Understanding the precise nature of these PMC-M1 interactions is of particular importance, as they appear to be implicated in the recovery of motor function following stroke (Frost et al. 2003; Dancause et al. 2005; Schulz et al. 2017; Paul et al. 2023).

PMC can exert modulatory influence on ipsilateral M1 output through direct cortico-cortical connections, refining movement parameters such as force and grip-type (Davare et al. 2008; Davare et al. 2009). Dual-site transcranial magnetic stimulation (TMS) studies have shown that PMd and PMv differentially modulate ipsilateral M1 excitability, with distinct patterns of inhibition and facilitation depending on the subregion stimulated and stimulus parameters (Davare et al. 2008; Groppa et al. 2012). These ipsilateral influences are supported by anatomical connectivity, with both PMd and PMv sending direct projections to ipsilateral M1 via short-range cortico-cortical fibers (Dancause et al. 2006; Schulz et al. 2014; Dea et al. 2016). However, the influence of PMC on M1 extends beyond the ipsilateral hemisphere to encompass interhemispheric interactions, which is the focus of the present study.

Anatomical evidence from tracing studies in non-human primates shows that PMC has many transcallosal connections with contralateral M1 (Marconi et al. 2003; Boussaoud et al. 2005; Dancause et al. 2007). In humans, diffusion-based tractography confirms the transcallosal representation of PMC within the corpus callosum and identifies PMd as a key contributor to the interhemispheric fiber tracts in the cortical motor network (Zarei et al. 2006; Ruddy et al. 2017). Neurophysiological evidence from dual-site TMS further supports this transcallosal connectivity, demonstrating that PMd can modulate contralateral M1 excitability across distinct interhemispheric inhibitory windows (Ni et al. 2009). Structural evidence also demonstrates interhemispheric connectivity between PMv and contralateral M1, though these connections appear less prominent than those from PMd (Boussaoud et al. 2005; Dancause et al. 2007; Ruddy et al. 2017). This notion is supported by dual-site TMS studies demonstrating complex PMv-to-M1 interhemispheric modulations (Fiori et al. 2017). Despite gathering anatomical and neurophysiological evidence for PMd-to-M1 and PMv-to-M1 interhemispheric connectivity, the precise nature of these influences — particularly how PMd and PMv differentially modulate contralateral M1 excitability — remains incompletely understood.

TMS offers complementary methods to probe the neurophysiology of interhemispheric inhibition and facilitation (Ferbert et al. 1992; Chen et al. 2008). Single-pulse TMS can elicit the ipsilateral silent period (iSP), a transient suppression of voluntary muscle activity in the contracting limb ipsilateral to the stimulated hemisphere, which reflects transcallosal inhibition (Ferbert et al. 1992; Hupfeld et al. 2020). The iSP provides a broad index of transcallosal inhibitory connectivity — including the silent period magnitude, onset, and duration — and is understood to be driven primarily by cortical mechanisms via the corpus callosum, though precise mediating mechanisms remain an area of investigation (Meyer et al. 1995; Boroojerdi et al. 1996; Hupfeld et al. 2020). Dual-site TMS offers a complementary approach that can further probe mechanisms of interhemispheric inhibition (IHI) and facilitation (IHF) (Ferbert et al. 1992; Chen et al. 2008; Neva et al. 2020). Dual-site TMS employs a conditioning stimulus (CS) over one hemisphere followed by a test stimulus (TS) over contralateral M1, allowing the examination of IHI/IHF across distinct temporal windows (Ferbert et al. 1992; Chen et al. 2008; Van Malderen et al. 2023). Two distinct temporal windows of M1-to-M1 IHI have been demonstrated, including short-IHI (ISIs of 8–12 ms) and long-IHI (ISIs of 30–50 ms; Ni et al. 2009). Each of these IHI phases reflect distinct underlying mechanisms of inhibition toward the contralateral M1, with evidence that long-IHI being mediated by postsynaptic GABA_B_ receptors, and short-IHI being less understood (Irlbacher et al. 2007). Importantly, long-IHI elicited from M1 has been shown to correlate with iSP duration, suggesting that these two measures capture overlapping transcallosal mechanisms of influence on contralateral M1 (Chen et al. 2003).

To date, the iSP has been employed exclusively to study M1-to-M1 interhemispheric communication, and its potential to assess transcallosal inhibition originating from non-M1 regions such as PMd and PMv has not been investigated (Neva et al. 2020). Dual-site TMS studies have demonstrated that a CS over PMd can modulate contralateral M1 excitability, eliciting short-IHI at ISIs of 6-10 ms (Mochizuki et al. 2004; Koch et al. 2006; Ni et al. 2009) and long-IHI at ISIs of 40-60 ms (Ni et al. 2009). For PMv, a CS has been shown to elicit long-IHI and facilitation at other long-latency ISIs (Fiori et al. 2017), but short-IHI mechanisms from PMv remain unexplored. Previous investigations of PMd-to-M1 and PMv-to-M1 interactions have been conducted in separate studies and under different stimulation parameters (e.g., ISIs), precluding direct comparison between the two subregions and with M1-to-M1 IHI within a single study (Mochizuki et al. 2004; Ni et al. 2009; Fiori et al. 2017). Importantly, at ISIs as short as 4-5 ms, contralateral M1 facilitation (IHF) can occur, which is thought to represent the minimum transcallosal conduction time in humans (Hanajima et al. 2001; Ni et al. 2020). Below this minimum conduction latency, any modulation of contralateral M1 output likely reflects non-callosal contributions (i.e., subcortical or spinal pathways), a notion supported by non-human primate evidence demonstrating modulation of contralateral M1 during simultaneous stimulation (Quessy et al. 2016; Côté et al. 2017). Critically, non-callosal IHI, as probed by a 0 ms ISI (i.e., simultaneous CS and TS delivery), has not been examined from PMd or PMv in humans, despite evidence for modulation of contralateral M1 output in non-human primates (Quessy et al. 2016; Côté et al. 2017).

No previous research has comprehensively examined the differential influence of PMd and PMv on contralateral M1 excitability, integrating both single-pulse (iSP) and dual-site TMS (short-, long-, and non-callosal IHI) within the same study. Therefore, the objective of this study was to characterize the differential interhemispheric influence of PMd and PMv on contralateral M1 output using two complementary TMS approaches. Here, we assessed the iSP to index overall transcallosal inhibition from PMd, PMv and M1, and employed dual-site TMS to examine short-(10 ms ISI), long-(50 ms ISI), and non-callosal (0 ms ISI) IHI mechanisms. We hypothesized that an iSP would be elicited from PMd and PMv with unique silent period parameters compared to each other and to M1. We also hypothesized that PMd and PMv would display distinct short-, long-and non-callosal-IHI profiles compared to each other and to M1 itself. Finally, based on previous findings from M1 (Chen et al. 2003), we expected that iSP duration would be uniquely associated with long-IHI when elicited from PMd and PMv.

## 2. MATERIALS AND METHODS

### Participants

Thirty right-handed adults (15 F, 18-39 years, mean age 26.4 [4.27]) took part in the study. Handedness was confirmed by using the Edinburgh Handedness Inventory (Oldfield 1971). A sensitivity analysis conducted using G*Power indicated that the current study had 90% power to detect an effect size f = 0.26 (equivalent to ∼*η²_p_* = 0.06), which corresponds to a medium effect size at an alpha level of.05 (Lakens 2022). Informed consent was obtained before participation in any experimental protocol. Participants were screened for any potential contraindications to TMS and magnetic resonance imaging (MRI) using standard screening forms prior to participation. Participants reported no neurological conditions or disorders. The experimental procedures for this study were approved by the Ethics Committee of the Centre de recherche de l’Institute Universitaire de Gériatrie de Montréal (CRIUGM) under the approval number of “CER VN 21-22-36”.

### Experimental design

This study used a within-subject design that included one preliminary visit and two experimental sessions involving different TMS assessments. The preliminary visit included acquisition of a T1-weighted structural MRI scan used for TMS targeting and position monitoring. The first experimental session included the assessment of transcallosal inhibition (TCI), via measurement of the iSP, elicited from right hemisphere M1, PMd and PMv. See *Experimental Session 1: iSP evaluation* for further details. The second experimental session included the use of dual-site TMS to assess the neurophysiological mechanisms of short-, long-and non-callosal-IHI as elicited from right hemisphere M1, PMd and PMv to left M1 with three distinct ISIs (0, 10 and 50 ms; Figure 1). See *Experimental Session 2: dual-site IHI evaluation* for further details. To accommodate diurnal fluctuations in M1 excitability (Merrell et al. 2024), the two experimental sessions were scheduled at the same time of day (±3 h) for each participant (Tamm et al. 2009).

**Figure 1.**
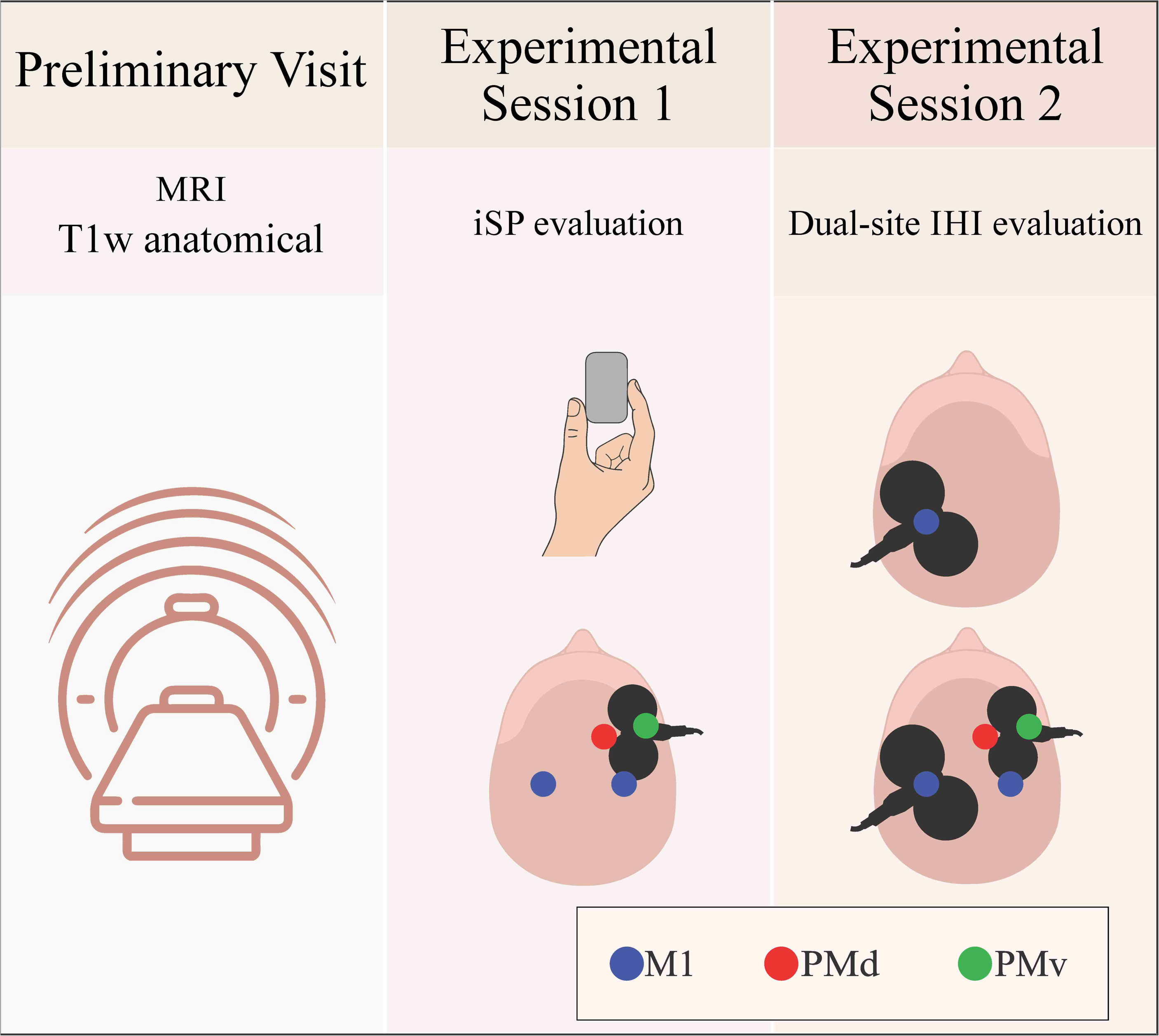
**Experimental design**. The study employed a within-subject design consisting of one preliminary visit and two experimental sessions. *Preliminary visit*: a T1w structural MRI scan was acquired for neuronavigation-guided transcranial magnetic stimulation (TMS) targeting and coil position monitoring. *Experimental Session 1*: transcallosal inhibition (TCI) was assessed via the ipsilateral silent period (iSP), elicited by suprathreshold single-pulse TMS over right hemisphere M1, PMd, and PMv while holding an isometric pinch grip contraction using the right hand (ipsilateral to the TMS pulse) at 50% of maximum voluntary contraction. *Experimental Session 2*: dual-site TMS evaluated short-, long-, and non-callosal-interhemispheric inhibition (IHI), with conditioning stimuli (CS) over right hemisphere M1, PMd, and PMv and the test stimulus (TS) over left M1, at three ISIs (0, 10, and 50 ms). Colored circles indicate TMS target locations: M1 (blue), PMd (red), and PMv (green). Both experimental sessions were conducted at the same time of day (±3 h).

### Electromyographic (EMG) recording

EMG data were collected with 1-cm diameter surface electrodes from first dorsal interosseous (FDI) muscle of both hands. The electrodes were placed over the FDI muscle in a belly-tendon configuration. The ground electrode was placed over the ulnar styloid process. EMG signals were collected using LabChart software (Labchart 8.0) with a PowerLab (PL3516 PowerLab, 16/35 16 Channel Recorder, AD Instruments, Colorado Springs, CO, USA) data acquisition system. EMG data was amplified by a bioamplifier (Dual Bio Amp, AD Instruments, Colorado Springs, CO, USA) with an acquisition rate of 2 kHz, bandpass (20-400 Hz) and notch filtered (center frequency of 50 Hz). Data were captured in a 500-ms sweep from 100 ms before to 400 ms after TMS delivery.

### Transcranial magnetic stimulation (TMS)

During the TMS assessments, participants were comfortably seated on an adjustable chair and remained at rest. Single-pulse and dual-site TMS were delivered with a Magstim BiStim^2^ stimulator (Magstim Co., UK) connected to a 7-cm outer diameter figure-of-eight coil (Magstim 70 mm P/N 9790, Magstim Co., UK) and a 5-cm outer diameter branding iron coil (Magstim Co., UK) over the left (dominant) and right (non-dominant) hemisphere M1 FDI representations, respectively. The TMS pulses generated were monophasic and with current flow in the posterior-to-anterior direction over left M1, and lateral-to-medial over right M1. The 7-cm outer diameter coil was oriented ∼45° to the mid-sagittal line with the handle facing posteriorly over left M1. The 5-cm outer diameter branding iron coil was oriented perpendicular to the mid-sagittal over right M1. To ensure accurate and consistent coil placement, we used individual participant anatomical MRI (T1-weighted) for real-time position monitoring and cortical region targeting using the BrainSight neuronavigation system (Rogue Research Inc., Montreal, QC, Canada). Positioning the respective coils over left and right M1, the “hotspot” for the M1 FDI representation was identified, which was determined as the cortical location over M1 that elicited the largest and most consistent MEPs in contralateral FDI. For the right M1 hotspot, resting motor threshold (RMT) was determined as the lowest stimulator output to elicit peak-to-peak motor evoked potential (MEPs) of at least 50 µV in 5 out of 10 consecutive trials (Rossini et al. 1994). During dual-site TMS, the stimulus intensity was adjusted to elicit consistent peak-to-peak MEP amplitudes of ∼1 mV from left and right M1, which were used for our test (TS) and conditioning stimulus (CS) intensities, respectively (Daskalakis et al. 2002). We defined PMd as 2 cm anterior and 1 cm medial relative to the right M1 hotspot (O’Shea et al. 2007; Ni et al. 2009), and PMv as 3 cm anterior and 2.5 cm lateral relative to the right M1 hotspot (Baumer et al. 2009). Additionally, using individual MRI neuronavigation, we ensured that the CS coil for PMd and PMv were over a gyrus rather than sulcus, slightly adjusting the CS coil placement accordingly (Ni et al. 2009). The Montreal Neurological Institute (MNI) coordinates for left M1, as well as right M1, PMd and PMv were recorded for each participant. These MNI coordinates were used for cortical target estimations, described further below.

### Experimental Session 1: iSP evaluation

Experimental session 1 investigated iSP via the use of single-pulse TMS over right M1, PMd and PMv. The iSP was measured while participants held an isometric contraction of 50% maximum voluntary contraction of the right FDI with a pincer grip using a hand-held dynamometer, which was connected to a PowerLab data acquisition system that provided real-time visual feedback using a custom program with LabChart software (Labchart 8.0). 20 single TMS pulses were delivered at 150% RMT using the 5-cm outer diameter branding iron coil (Magstim Co., UK) over right M1, PMd and PMv individually (3 regions, 20 pulses / region = 60 pulses total) (Jung and Ziemann 2006; Neva et al. 2016). Participants were given rest periods of 30-60 s after collection of 10 trials with the purpose of avoiding fatigue.

### Experimental Session 2: Dual-site IHI evaluation

In Experimental session 2, we further investigated the underlying mechanisms of IHI as assessed with dual-site TMS. The stimulation intensity required to elicit a ∼1 mV MEP from the left M1 hotspot was used as the TS intensity and the intensity required to elicit ∼1 mV MEP from the right M1 hotspot was applied to right PMd and PMv as the CS intensity. Dual-site IHI assessment consisted of delivering a CS over right PMd/PMv/M1 followed by a TS over left M1 at different ISIs (Ni et al. 2009; Neva et al. 2015; Fiori et al. 2017). Three distinct ISIs between CS and TS were evaluated: 0, 10 and 50 ms. These ISIs were chosen to investigate the influence of PMd, PMv and M1 on contralateral M1 excitable output via short (10 ms)-, long (50 ms)-, and non-callosal (0 ms)-IHI mechanisms (Ni et al. 2009; Côté et al. 2017). During IHI assessments elicited from PMd and PMv, we ensured that MEPs were not elicited from the CS in the contralateral FDI. To do this, when we observed MEPs from contralateral FDI, we systematically decreased the CS intensity in steps of 2% MSO until there were no observable MEPs. For PMd, one single participant displayed MEPs in contralateral FDI and the %MSO was decreased by a total of 6% MSO. No MEPs were observed in the contralateral FDI when applying the CS over PMv. Ten CS+TS trials and 10 TS alone trials were recorded for each cortical region (PMd, PMv and M1) and each ISI (0, 10 and 50 ms; 90 CS+TS pulses and 90 TS pulses = 180 total pulses).

### Data processing and statistical analysis

Repeated measures analysis of variance (RM-ANOVA) was used to analyze the data from Experimental session 1 (iSP data) and 2 (dual-site IHI data). The specific analyses will be described below in more detail. Residual statistics, skewness, and kurtosis values, along with plots, were generated to assess the normality and homoscedasticity of the data. Normality of data was further evaluated using the Shapiro-Wilk test with a significance level set at p <.001 (Gamst 2008). To satisfy the normality assumption, the Greenhouse-Geisser correction was applied when normality was violated for RM-ANOVAs (variables: M1 0 ms and PMd 0 ms: W_30_ ≥.683, ps ≥.001) and data were square root-transformed for paired t-tests (variables: M1 at 0 and 10 ms, PMd at 0 ms, PMv at 0,10 and 50 ms; W_30_ ≥.639, ps ≥.001). Untransformed data are displayed in figures. *Post hoc* analyses were performed using Tukey’s HSD as appropriate. Effect sizes were calculated and reported as partial eta squared (*η²_p_*) following established guidelines for interpretation (Cohen 2013) where 0.01 indicates a small effect, 0.06 a moderate effect, and 0.14 a large effect. Statistical procedures were conducted by SPSS (SPSS 28.0) software. All data are reported as mean (SD) unless otherwise stated. Statistical significance was set at p <.05.

*EMG data processing for MEP amplitudes, iSP and IHI.* For iSP and IHI assessment, we visually inspected all EMG data to screen for voluntary muscle activity. Any trials showing visible voluntary pre-stimulus EMG activity from the non-dominant (resting) FDI during iSP assessment and bilateral FDI during IHI assessment were excluded from analysis (constituting 0.52% of trials). Specific data processing and analysis for Experimental sessions 1 and 2 are detailed further below.

*Baseline neurophysiological data: TS %MSO and TS MEP amplitudes*. Stable TS %MSO values during IHI assessment across cortical regions were ensured using a one-way RM-ANOVA, considering within-subject factor REGION (PMd, PMv, M1). Stable TS MEP amplitudes during IHI assessment across cortical regions were also ensured using one-way RM-ANOVA, considering within-subject factor REGION (M1, PMd, PMv).

*MNI Coordinate Estimation.* To estimate the anatomical location of premotor TMS targets, group-average MNI coordinates were analyzed using two complementary approaches. First, the SPM Anatomy Toolbox v3.0 (Eickhoff et al. 2005) was used to assess localization to cortical gyri and Brodmann areas based on probabilistic cytoarchitectonic maps. Second, Neurosynth (Yarkoni et al. 2011), a meta-analytic database of over 14,000 fMRI studies, was used to evaluate functional associations with premotor terms. Additionally, anatomical separation between targets was quantified by calculating Euclidean distances in MNI space. Specifically, PMd-to-M1 distance was calculated to assess spatial distinctness from M1 representation, and vertical (superior-inferior) separation between PMd and PMv was calculated to assess distinction between dorsal and ventral premotor areas. Recorded z-coordinates were reduced by 20 mm to account for scalp-to-cortex distance prior to anatomical analysis, consistent with previous work (Opitz et al. 2011).

*Experimental Session 1*. Transcallosal inhibition elicited from the premotor cortices (PMd and PMv) and M1 was quantified by the iSP. The iSP was defined as the transient reduction in volitional EMG activity elicited by TMS applied over the cortex ipsilateral to the active muscle (Fling and Seidler 2012; Borich et al. 2015). To calculate the iSP as elicited from each region (PMd, PMv and M1), EMG data full were wave rectified and averaged for each participant. Pre-stimulus mean EMG (pre-stimulus_mean_) was defined as the voluntary EMG signal recorded 100 ms prior to TMS delivery. The iSP onset (iSP_onset_) was defined as the post-TMS time point (in ms) where the rectified EMG fell below the pre-stimulus_mean_ EMG signal and continued to decrease to less than 2 standard deviations below this level. The iSP offset (iSP_offset_) was defined as the point at which the EMG signal resumed the level of the pre-stimulus_mean_ activity consistently for a minimum of 2 ms (Jung and Ziemann 2006; Fling and Seidler 2012). The iSP duration comprised all data points between iSP_onset_ and iSP_offset_ (calculated in ms). The average rectified EMG level during the iSP was defined as the iSP mean (iSP_mean_) (Jung and Ziemann 2006). The iSP_mean_ was then expressed as a ratio of the mean pre-stimulus EMG (iSP_mean_/pre-stimulus_mean_), where a lower value indicates more inhibition. Custom MATLAB scripts (Mathworks, Natick, MA) were used to identify parameters of the iSP for each participant (Mang et al. 2015; Neva et al. 2016). While collecting the iSP elicited from PMd, we observed the appearance of contralateral MEPs. We thus quantified the mean peak-to-peak MEP amplitudes from PMd and compared them to those elicited from M1. No MEPs were elicited during iSP elicited from PMv.

Our first analysis quantified whether a significant iSP was elicited following stimulation over PMd, PMv and M1. To do this, we compared the iSP_mean_ to the pre-stimulus_mean_ EMG using a two-way RM-ANOVA with within-subjects factors EMG-SIGNAL (iSP_mean_, pre-stimulus_mean_) and REGION (PMd, PMv, M1). Our second analysis further quantified iSP parameters (iSP_mean_/pre-stimulus_mean_, iSP_onset_ and iSP duration), performing one-way RM-ANOVAs using the within-subjects factor REGION (PMd, PMv, M1). Finally, paired t-tests were performed comparing the mean peak-to-peak MEP amplitudes elicited from PMd and M1 during iSP collection.

*Experimental Session 2*. Dual-site paired-pulse TMS was used to assess short-, long-and potential non-callosal mechanisms of IHI from the right premotor cortices and M1 to left M1. Custom MATLAB scripts were used to assess peak-to-peak MEP amplitudes (measured in millivolts, mV) as elicited by TS alone and CS+TS for each cortical region. IHI magnitude was quantified as a ratio of CS + TS MEP amplitude over TS MEP amplitude (Kujirai et al. 1993): IHI ratio = (CS+TS)/TS, where smaller values indicate greater inhibition, while larger values indicate less inhibition (disinhibition or facilitation). This was performed for each cortical region (PMd, PMv, M1) and ISI (0, 10, and 50 ms). The ISIs were specifically chosen to investigate short (10 ms)-and long (50 ms)-, and potential non-callosal (0 ms)-IHI mechanisms impacting contralateral M1 output (Mochizuki et al. 2004; Ni et al. 2009; Côté et al. 2017; Fiori et al. 2017).

Our first analysis assessed inhibition of left M1 output elicited by a CS over right hemisphere PMd, PMv or M1. To do this, we compared peak-to-peak MEP amplitudes elicited with TS alone over left M1 to those elicited from CS+TS. Paired t-tests compared MEP amplitude elicited with a TS alone to CS+TS within each cortical region across ISIs (10, 50 and 0 ms; Bonferroni correction applied as appropriate; Ni et al. 2009). Our second analysis assessed IHI magnitude between cortical regions and ISIs using a two-way RM-ANOVA involving within-subjects factors REGION (PMd, PMv, M1) and ISI (0, 10, 50 ms), using IHI ratio as the dependent measure.

*Correlational analyses of Experimental Session 1 (iSP) and 2 (IHI) data*. Bivariate correlational analyses (Pearson’s r_p_ for normally distributed data or Spearman’s r_s_ for not normally distributed data) were conducted to characterize the strength in relationships between iSP duration and long-IHI (i.e., 50 ms ISI). As previous research has shown that M1 iSP duration was associated with M1 IHI at longer ISIs (e.g., 40 ms) but not shorter ISIs (Chen et al. 2003), we examined whether PMd/PMv demonstrated similar associations.

## 3. RESULTS

### Baseline neurophysiological data

For TS %MSO values during IHI assessments, there was no significant main effect of REGION (F_2, 58_ = 0.222, p =.802, *η²_p_* =.008), confirming stable TS intensity during assessment of IHI across the different brain regions examined. For TS MEP amplitudes during IHI assessments, there was no significant main effect of REGION (F_2, 58_ =.125, p =.882, *η²_p_* =.004), confirming consistent corticospinal output excitability throughout the assessment of IHI across the different brain regions (see *Supplementary Table 1*).

### MNI Coordinate Estimation

Group-average MNI coordinates (x, y, z) for the TMS targets were: right M1 (47.5,-10.2, 59.5), right PMd (40.4, 11.2, 57.7), and right PMv (61.8, 12.4, 28.5). SPM Anatomy Toolbox analysis localized the PMd coordinates to the middle frontal gyrus (macroanatomy: 66%), positioned immediately anterior to the cytoarchitectonic border of Area 6. The PMv coordinates were localized to the ventral precentral gyrus / Area 44 transition zone (cytoarchitecture: Area 44, 15.2%; macroanatomy: precentral gyrus, 31%). Neurosynth functional decoding revealed strong associations with premotor function for both targets (PMd: posterior probability = 0.60 for “dorsal premotor cortex”; PMv: posterior probability = 0.56 for “ventral premotor”). Euclidean distance between the PMd target and the M1 hotspot was 22.6 mm, with the primary separation along the anterior-posterior axis (Δy = 21.4 mm). The vertical separation between PMd and PMv targets was 32.7 mm (Δz), suggesting distinct dorsal and ventral premotor localizations.

### Experimental Session 1: Ipsilateral silent period (iSP) elicited from PMd, PMv and M1

Figure 2 displays representative data of individual EMG traces, mean and individual data demonstrating iSPs elicited from M1, PMd and PMv. Two-way RM-ANOVA revealed a significant interaction between EMG-SIGNAL and REGION (F_2, 58_ = 9.264), p <.001, *η²_p_* =.242; Figure 2A-B). Post hoc analyses demonstrated a significant decrease in rectified EMG during the iSP_mean_ compared to the pre-stimulus_mean_ for PMd (p <.001), PMv (p <.001) and M1 (p <.001; Figure 2). Main effects of EMG-SIGNAL (F_1, 29_ = 110.560, p <.001, η*²_p_* =.792) and REGION (F_2, 58_ = 6.043, p =.004, *η²_p_* =.172; Figure 2A-B) were also found.

**Figure 2.**
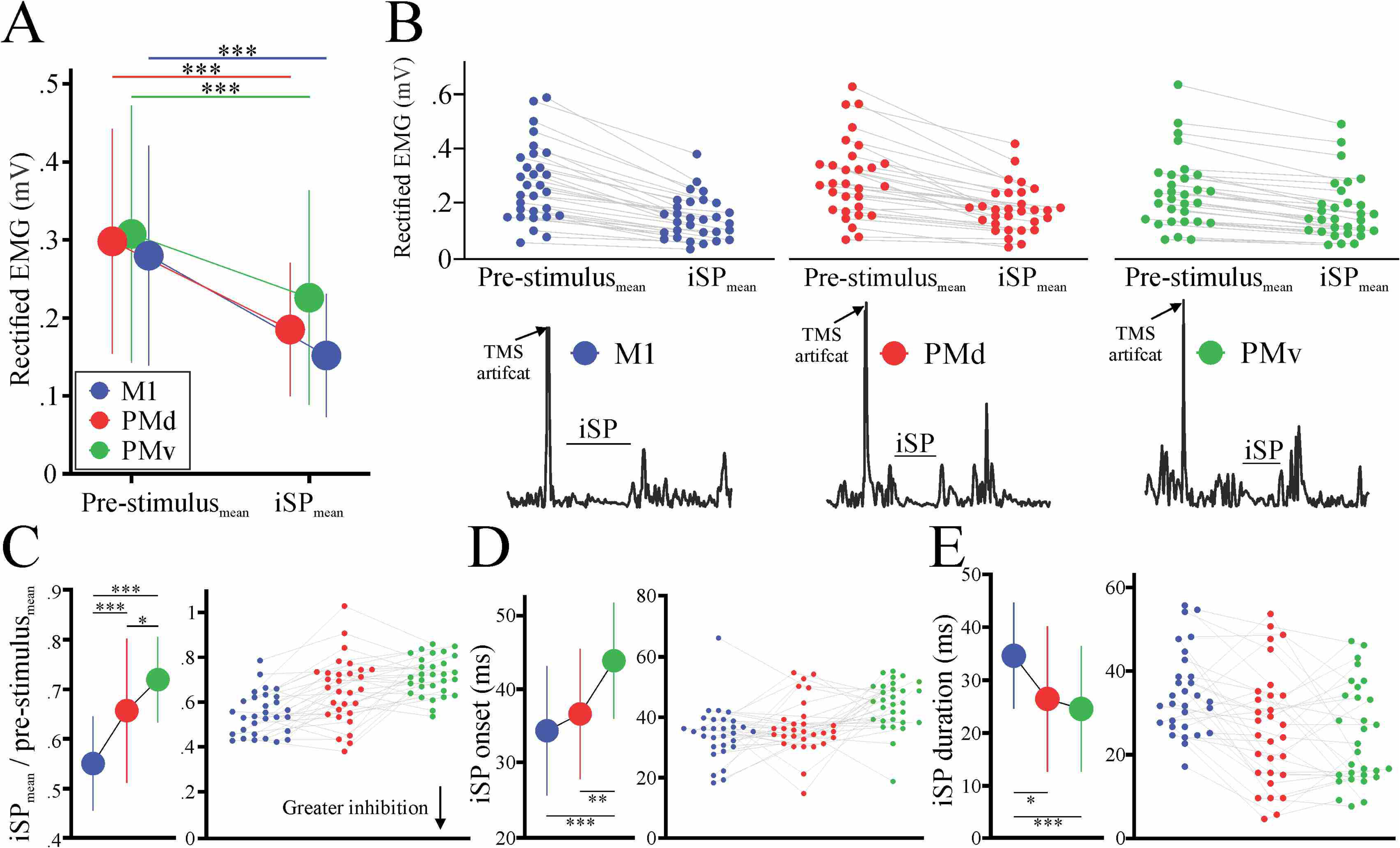
Experimental Session 1: iSP results. (A) Group mean (± SD) rectified EMG amplitude (mV) for the pre-stimulus period and iSP window across stimulation regions (M1, blue; PMd, red; PMv, green). (B) *Top*: Individual participant data showing rectified EMG amplitude during the pre-stimulus period and iSP window for M1, PMd, and PMv, with grey lines connecting paired observations. *Bottom*: Representative single-participant EMG traces illustrating the iSP following suprathreshold TMS over M1, PMd, and PMv, with the TMS artifact and iSP window indicated. (C) iSP magnitude, expressed as the ratio of mean EMG during the iSP to pre-stimulus mean EMG (iSP_mean_/pre-stimulus_mean_), shown as group means ± SD (left panel) and individual participant data with connected observations (right panel). Lower values indicate greater inhibition. (D) iSP onset latency (ms) shown as group means ± SD (left panel) and individual participant data (right panel). (E) iSP duration (ms) shown as group means ± SD (left panel) and individual participant data (right panel). In all panels, asterisks denote significant post hoc differences (* p <.05, ** p <.01, *** p <.001).

For iSP_mean_/pre-stimulus_mean_ one-way RM-ANOVA revealed a significant main effect of REGION (F_2, 58_ = 29.127, p <.001, *η²_p_* =.501; Figure 2C), with post hoc analyses showing that M1 (p <.001) displayed the greatest magnitude of inhibition, followed by PMd (p <.001), then PMv (p = 0.01). For iSP_onset_, one-way RM-ANOVA revealed a significant main effect of REGION (F_2, 58_ = 10.229, p <.001, *η²_p_* =.261; Figure 2D), with post hoc analyses showing both M1 (p <.001) and PMd (p =.004) displayed shorter iSP onsets compared to PMv, with no difference between M1 and PMd (p =.550). Finally, for iSP duration, one-way RM-ANOVA revealed a significant main effect of REGION (F_2, 58_ = 5.976, p =.004, *η²_p_* = 0.121; Figure 2E), with M1 displaying significantly longer duration than PMd (p =.026) and PMv (p=.005), while PMd and PMv demonstrated similar durations (p =.819).

*Assessment of MEPs elicited in contralateral (left) FDI during iSP*. A paired t-test demonstrated that the peak-to-peak MEP amplitude elicited when assessing iSPs from PMd and M1 to be significantly different (p <.001). Specifically, peak-to-peak MEP amplitudes elicited from M1 (5.1 ± 2.3 mV) were significantly greater than those elicited from PMd (2.9 ± 2.1 mV). No MEPs were observed when assessing iSP elicited from PMv.

### Experimental Session 2: Dual-site IHI assessment from PMd, PMv and M1

Figure 3 displays representative data of individual EMG traces, mean and individual data of MEPs elicited from TS alone and CS+TS as well as IHI ratio data from M1, PMd and PMv for each ISI. For the 10 ms ISI, paired t-tests showed significant decrease in MEP amplitude as elicited from the CS+TS compared to TS alone for PMd (p <.001) and M1 (p =.001), but not PMv (p =.886; Figure 3B). For the 50 ms ISI, paired t-tests showed MEP amplitude decrease as elicited from CS+TS compared to TS alone for PMd (p <.001), M1 (p =.011) and PMv (p =.007; Figure 3C). Finally, for the 0 ms ISI, paired t-tests showed significant reductions in MEP amplitude as elicited from CS+TS compared to TS alone for PMd, PMv and M1 (ps <.001; Figure 3A).

**Figure 3.**
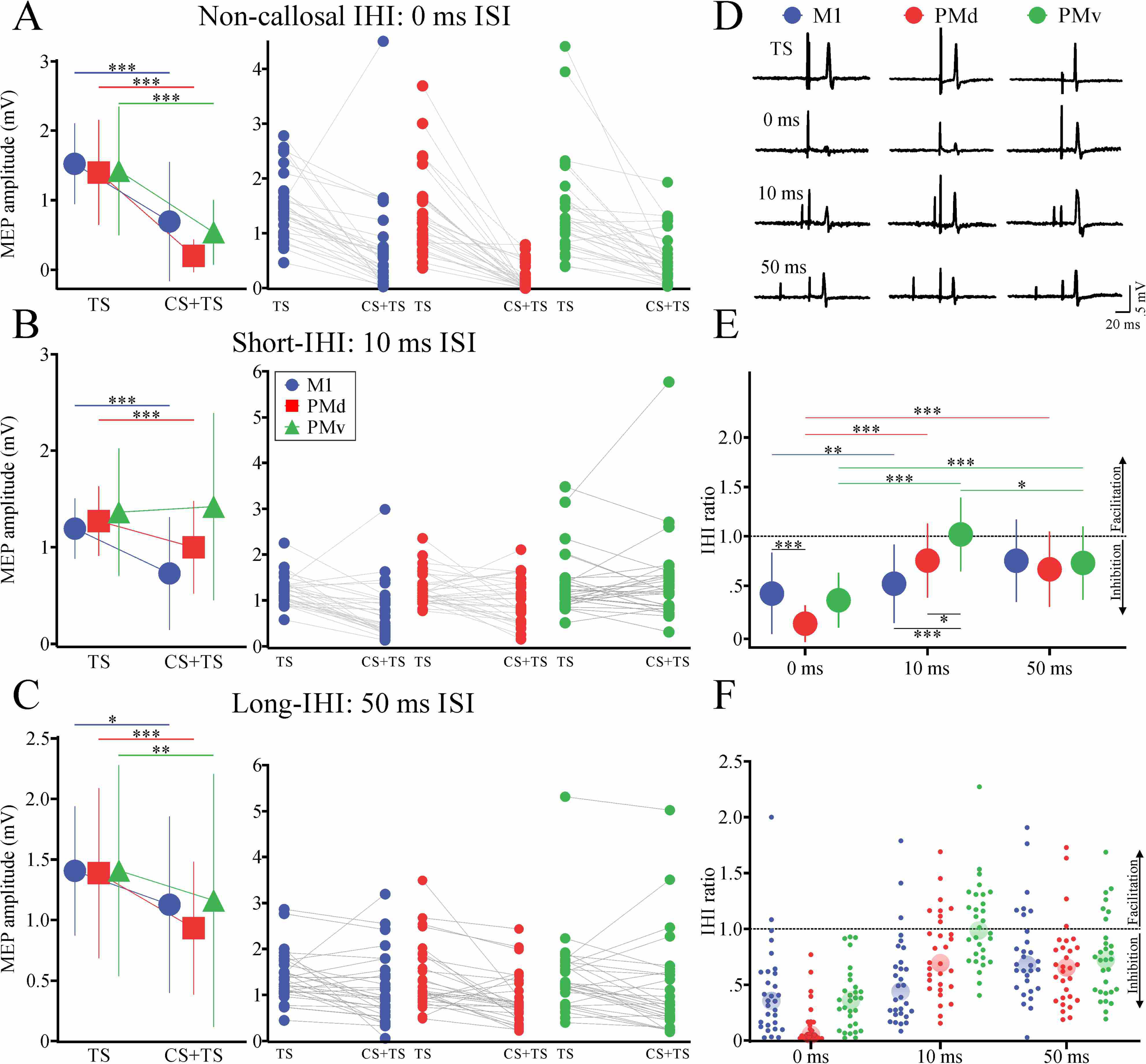
Experimental Session 2: IHI results. (A–C) *Left panels*: Group mean (± SD) MEP amplitude (mV) for the test stimulus alone (TS) and the conditioned stimulus paired with the test stimulus (CS+TS) for M1 (blue circles), PMd (red squares), and PMv (green triangles), at ISIs of 0 ms (A: Non-callsoal IHI), 10 ms (B: Short-IHI), and 50 ms (C: Long-IHI). *Right panels*: Individual participant MEP amplitudes for TS and CS+TS across all three stimulation sites at each ISI, with grey lines connecting paired observations. (D) Representative single-participant EMG traces for TS alone and CS+TS conditions at each ISI (0, 10, and 50 ms), shown separately for each conditioning site (M1, PMd, PMv). (E) Group mean (± SD) and (F) individual participant IHI ratios (CS+TS MEP / TS MEP) across ISIs for M1, PMd, and PMv. The dashed horizontal line at 1.0 demarcates inhibition (below) from facilitation (above). In all panels, asterisks denote significant post hoc differences (* p <.05, ** p <.01, *** p <.001).

Figure 3E-F displays mean and individual IHI ratio data. Two-way RM-ANOVA revealed a significant interaction between REGION and ISI (F_3.34, 96.74_= 8.895, p <.001, *η²_p_* =.235). For the 10 ms ISI, post hoc analyses demonstrated that M1 (p <.001) and PMd (p =.048) displayed greater inhibition than PMv, with no difference between M1 and PMd (p =.119). For the 50 ms ISI, there were no differences between cortical regions (ps >.980). For the 0 ms ISI, post hoc analyses demonstrated that PMd displayed greater inhibition compared to M1 (p =.001), with no other differences (ps >.955). Within PMd, post hoc analyses demonstrated greater inhibition with a 0 ms compared to both 10 (p <.001) and 50 ms (p <.001) ISIs, with no difference between 10 and 50 ms (p =.977) ISIs. Within PMv, post hoc analyses demonstrated greater inhibition with a 0 ms ISI compared to both 10 and 50 ms ISIs (ps <.001), along with greater inhibition elicited with a 50 ms compared to 10 ms ISI (p =.021). Within M1, post hoc analyses demonstrated greater inhibition elicited with a 0 ms compared to a 50 ms (p =.003) ISI, with no other differences (ps >.125). We also observed main effects of ISI (F_1.99, 57.57_ = 48.134, p <.001, *η²_p_* =.624) and REGION (F_1.79, 51.88_= 6.812, p =.002, *η²_p_* =.190).

### Correlational analyses between Experimental Session 1 (iSP) and 2 (IHI ratio) data

Figure 4 displays correlational analysis between iSP duration and long-IHI tested with a 50 ms ISI for each region. We found a significant negative correlation for PMd (r_s_ = -.404, R² =.163, p =.027) and PMv (r_s_ = -.365, R² =.133, p =.047), while we did not for M1 (r_s_ = -.021, R² =.0004, p =.912).

**Figure 4.**
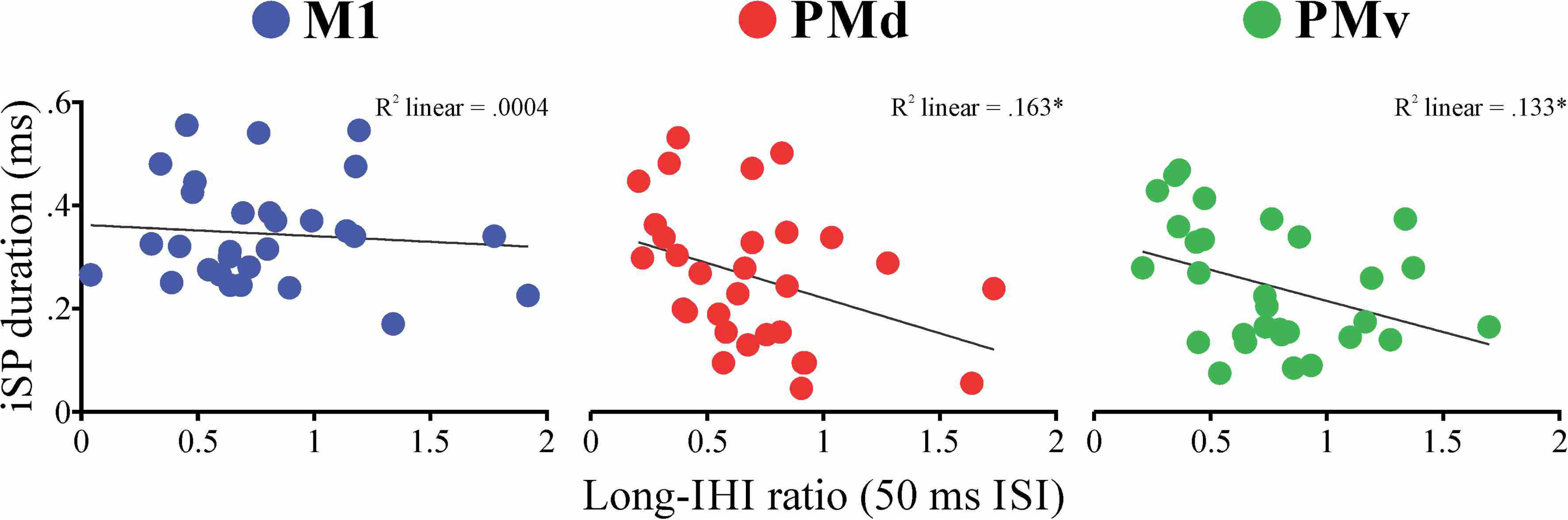
Correlations between iSP duration and long-IHI ratio. Scatterplots illustrating the relationship between iSP duration (ms; Experimental Session 1) and the long-IHI ratio (50 ms ISI; Experimental Session 2) for M1 (blue, *left*), PMd (red, *middle*), and PMv (green, *right*). Each data point represents an individual participant. The linear line of best fit and corresponding R² value are displayed in each panel. Asterisks (*) denote statistically significant correlations (p <.05).

## 4. DISCUSSION

This study provides a systematic characterization of how PMd and PMv influence contralateral M1 output, revealing that these PMC subregions display distinct modulation profiles compared to each other and to M1. Our study highlights two primary novel findings: 1) an iSP can be elicited from PMd and PMv, with each premotor region displaying unique silent period properties compared to each other and to M1; 2) non-callosal IHI was demonstrated at the 0 ms ISI, revealing that PMd produces stronger, and PMv elicits similar inhibition at this interval compared to M1. We also found that PMd and PMv displayed greater diversity in their inhibitory profiles across short-, long-and non-callosal-IHI in comparison to M1. Finally, iSP duration correlated with long-IHI magnitude for PMd and PMv, linking these two measures and suggesting they reflect partly overlapping neural mechanisms. Collectively, these findings position premotor cortices as dynamic contributors to interhemispheric interactions with M1, complementing our understanding of M1-mediated pathways. Here, we will discuss the potential explanations for, and mechanisms supporting, our findings.

### PMd and PMv elicit distinct ipsilateral silent periods

To our knowledge, the iSP has only been elicited from M1 in healthy and clinical populations. Here, we demonstrated that PMd and PMv also elicit iSPs, with largely unique silent period properties distinct from M1 and from each other (see *Results* for details). Our demonstration of an iSP elicited from PMd and PMv aligns with previous dual-site TMS studies showing IHI from these premotor regions to contralateral M1 (Mochizuki et al. 2004; Ni et al. 2009; Fiori et al. 2017), suggesting both techniques engage shared transcallosal inhibitory pathways, albeit across different temporal windows. Our findings extend earlier dual-site TMS work which showed that PMd and PMv exhibit distinct inhibitory profiles compared to M1 (Ni et al. 2009; Fiori et al. 2017). Notably, Ni et al. (2009) found that PMd-elicited IHI emerged at fewer interstimulus intervals compared to M1-elicited IHI, with a slight shift towards longer intervals in long-latency IHI (Ni et al. 2009). Our iSP results parallel these observations: PMd showed reduced iSP magnitude compared to M1, consistent with comparatively less engagement of transcallosal inhibitory pathways. PMd onset latencies, however, were comparable to M1, suggesting that while inhibitory strength differs, transcallosal conduction timing between the two regions is similar. These parallels are meaningful given that iSP likely reflects long-IHI mechanisms (Chen et al. 2003), and Ni et al. demonstrated that PMd-elicited IHI was prominent at longer-latency ISIs (≥40 ms) (Ni et al. 2009). The longer iSP onset latency for PMv compared to M1 and PMd likely reflects its distinct terminal organization of their corticospinal projections. PMv compared to PMd has fewer direct projections to the lower cervical enlargement, potentially requiring more complex and indirect pathways (Borra et al. 2010). In contrast, PMd has more robust (∼5 times) corticospinal terminal projection to cervical enlargement (Morecraft et al. 2019). These differences between PMd and PMv can explain shorter iSP onset latency from PMd. Together, these findings suggest that PMd engages transcallosal pathways with distinct temporal dynamics compared to M1, which can be assessed via the iSP using single-pulse TMS.

A critical methodological consideration in this study is whether the iSP elicited from premotor regions truly reflects PMC-to-M1 inhibition or from the current spread/indirect activation of ipsilateral M1. Several lines of evidence from our data support the notion that the iSPs elicited from PMd, PMv, and M1 each arise from anatomically and functionally distinct cortical sources. First, although significant reductions in ongoing EMG activity were observed from all three cortical stimulation sites, the iSP magnitude via normalized EMG followed a systematic gradient (M1 > PMd > PMv). If the PMd or PMv results were merely due to current spread to or indirect activation of ipsilateral M1, we would expect the elicited inhibition to more closely resemble the M1 iSP in magnitude, onset and duration. Instead, PMd and PMv displayed unique iSP durations and the onset latency elicited from PMv was distinct from M1; while PMd onset latency was not statistically different, there was a slight delay compared to M1. Second, our correlational analyses revealed region-specific associations. We found that the iSP duration elicited from PMd was significantly correlated with the magnitude of long-latency IHI from PMd. The same correlational pattern was found for PMv. Crucially, these relationships were specific to PMd and PMv, with M1 not displaying this correlational pattern with long-IHI, in contrast to previous findings (Chen et al. 2003). Nevertheless, our findings suggests that the inhibitory circuits activated by single-pulse TMS over PMd and PMv are distinct, at least in part, from those activated over M1. Third, MNI coordinate analysis demonstrated anatomical distinctness between stimulation sites. The PMd target was located 22.6 mm from the M1 hotspot, placing it in the middle frontal gyrus at the anterior border of Area 6 (Picard and Strick 2001; Chouinard and Paus 2006). This separation exceeds the estimated spread of TMS electric fields previously reported (∼10-15 mm; Thielscher and Kammer 2004), suggesting that PMd stimulation likely did not directly activate the M1 hand representation. Furthermore, PMd and PMv were separated by 32.7 mm vertically (superior-inferior axis), with PMv localized to the ventral precentral gyrus / Area 44 transition zone, supporting anatomical separation between premotor regions (Picard and Strick 2001; Chouinard and Paus 2006). Collectively, these findings provide evidence that the iSP can be used as a marker of PMC-to-M1 transcallosal inhibition, allowing the further distinction between interhemispheric connectivity from PMd and PMv to contralateral M1.

### PMd and PMv exhibit diverse interhemispheric inhibitory profiles

Our dual-site TMS results assessed the inhibitory profiles of PMd and PMv across different temporal windows, revealing that these premotor regions exerted differential influences on contralateral M1, while also having distinct patterns to M1-to-M1 IHI.

IHI elicited from PMd at short (10 ms) and long (50 ms) ISIs likely reflects distinct transcallosal cortico-cortical pathways (Ferbert et al. 1992; Irlbacher et al. 2007; Ni et al. 2009). The timing of short-IHI results is consistent with relatively fast transcallosal transmission, suggesting PMd projects to contralateral M1 either directly or via contralateral PMd. Previous studies demonstrated PMd-elicited short-IHI at 6, 8, and 10 ms ISIs (Mochizuki et al. 2004; Koch et al. 2006; Ni et al. 2009). While the specific pathway mediating short-IHI remains uncertain, M1-to-M1 SIHI (6-12 ms ISIs) is thought to involve transcallosal signals interacting with intracortical circuitry in contralateral M1, potentially through mechanisms of short-interval intracortical inhibition (Daskalakis et al. 2002). Given that PMd and M1 produced similar degrees of short-IHI in our study, PMd-elicited inhibition may engage similar intracortical mechanisms in contralateral M1, though this needs to be tested in future work. Long-IHI (50 ms) from PMd aligns with previous findings showing that PMd modulates contralateral M1 output through slower, potentially GABA_B_-mediated intracortical circuits at ISIs of 40-60 ms (Ni et al. 2009). M1-elicited long-IHI (30-50 ms ISIs) is believed to involve polysynaptic circuits or additional intracortical loops beyond that of short-IHI (Udupa et al. 2010; Ghosh et al. 2013). It is possible that PMd-to-M1 and M1-to-M1 long-IHI share similar mechanisms due to the similar magnitude and relative timing of inhibition as shown with our dual-site TMS results. However, the slight shift toward longer intervals in long-latency IHI elicited from PMd compared to M1 found in previous work (Ni et al. 2009), along with differences in our iSP parameters elicited from PMd compared to M1, suggests potentially distinct pathways of connectivity between PMd and contralateral M1 compared to M1-to-M1 interactions; yet this warrants future investigation.

A striking novel finding was the robust inhibition elicited from PMd at the 0 ms ISI. Given that 0 ms falls below the minimum conduction time for transcallosal communication in humans (∼4-5 ms; Hanajima et al. 2001; Ni et al. 2020), this effect is likely not mediated by cortico-cortical pathways. Instead, this non-callosal IHI likely reflects convergence of descending volleys from PMd and contralateral M1 at a subcortical or spinal level. This finding contrasts with previous animal studies showing that simultaneous stimulation of PMd and contralateral M1 produces predominantly facilitation (Quessy et al. 2016; Côté et al. 2017). Two methodological differences may explain this discrepancy. First, indwelling microelectrodes in non-human primates produce focal activation, likely recruiting fewer inhibitory interneurons and favoring facilitatory pathways, compared to TMS in humans (Thielscher and Kammer 2004; Tian and Izumi 2022). Second, we used relatively higher conditioning stimulus intensities (∼1 mV MEP) compared to animal studies (75% of EMG threshold in the contralateral arm to the conditioning electrode; Côté et al. 2017).

Our approach thus potentially recruited larger populations of inhibitory intracortical circuits, whose excitatory projections may preferentially synapse onto inhibitory interneurons at convergence sites with contralateral M1 volleys, resulting in a net suppression of M1 output. Future studies should systematically vary CS and TS intensities at 0 ms ISI to further investigate the potential interaction between inhibitory and facilitatory mechanisms contributing to this inhibition of contralateral M1 output. The reticular formation represents a plausible convergence site, given anatomical evidence that PMd sends dense corticoreticular projections to the pontomedullary reticular formation in animal models — more so than M1 — supporting a preferential PMd-reticular formation pathways (Keizer and Kuypers 1989; Kably and Drew 1998; Fregosi et al. 2017). Notably, PMd elicited significantly greater non-callosal IHI than M1 itself at 0 ms ISI, which is consistent with the dense corticoreticular connectivity of the premotor cortex more broadly (Yeo et al. 2012; Jang and Lee 2019), and with anatomical evidence of dense PMd projections to the pontomedullary reticular formation relative to M1 (Fregosi et al. 2017). We also cannot discount the fact that these descending signals may interact with other brainstem structures or at the spinal level before reaching the target muscle, as previous work has noted (Côté et al. 2017). Further investigation in humans and animal models is needed to uncover the pathways and mechanisms underlying PMd-elicited non-callosal IHI.

In contrast to PMd, PMv displayed a distinct inhibitory profile characterized by absent short-IHI and preserved long-IHI. PMv did not elicit significant inhibition at the 10 ms ISI, differing from both PMd and M1. However, PMv produced long-IHI at 50 ms ISI, comparable in magnitude to PMd and M1. These findings align with Fiori et al. (2017), who demonstrated PMv-elicited long-latency IHI at 40 ms and 150 ms ISIs (Fiori et al. 2017). The absence of short-IHI combined with preserved long-IHI suggests that PMv influences contralateral M1 through slower, potentially polysynaptic pathways involving higher-level cortical (e.g., prefrontal and parietal cortices) or subcortical relays (e.g., thalamus, brainstem structures) rather than direct transcallosal projections (Fiori et al. 2017; Ruddy et al. 2017). Our iSP results contribute evidence to this unique temporal signaling, where PMv showed significantly longer onset latency and reduced magnitude compared to PMd and M1. Our distinct iSP findings between premotor regions suggest that while PMv engages transcallosal pathways, these connections may be functionally distinct or less direct than those from PMd (Fiori et al. 2017; Ruddy et al. 2017). Importantly, the absence of short-IHI in our study does not rule out direct transcallosal PMv-to-M1 connectivity. Anatomical evidence confirms direct PMv projections to contralateral M1, particularly in rostral M1 where there is less digit representation (Dancause et al. 2007). Further, the difference in callosal fiber density from PMv and PMd to contralateral M1 is relatively small (Boussaoud et al. 2005), suggesting the functional distinctions observed here may reflect differences in pathway efficacy rather than anatomy alone. Collectively, these findings suggest that PMv-to-M1 interhemispheric communication operates through physiologically distinct mechanisms compared to PMd and M1 (Ruddy et al. 2017). Future research combining neurophysiological and structural connectivity approaches would be beneficial to clarify these potential pathway differences.

PMv produced robust non-callosal IHI at 0 ms ISI comparable in magnitude to PMd and M1. Our findings align with previous non-human primate work showing that simultaneous microstimulation of PMv and contralateral M1 primarily elicited inhibition of M1 output (Quessy et al. 2016; Côté et al. 2017). As with PMd, this effect is likely mediated by convergence of descending volleys at subcortical sites (e.g., reticular formation, spinal cord) rather than transcallosal pathways (see section on non-callosal IHI elicited from PMd above). Anatomical studies show that PMv also projects to the pontomedullary reticular formation and medulla, though with different density and somatotopic targets compared to PMd (Kably and Drew 1998; Wise 2006), suggesting a shared but potentially anatomically distinct subcortical mechanism.

### M1-to-M1 non-callosal interhemispheric inhibition

Our finding of robust inhibition with simultaneous bilateral M1 stimulation (0 ms ISI) is, to our knowledge, novel. Previous work has characterized distinct temporal phases of M1-to-M1 modulation: long-IHI at ∼40-60 ms ISIs (Ni et al. 2009), short-IHI at ∼10 ms ISIs (Hanajima et al. 2001; Daskalakis et al. 2002; Ni et al. 2009; Ni et al. 2020), facilitation at 4-5 ms ISIs likely reflecting initial excitatory transcallosal transmission (Hanajima et al. 2001; Ni et al. 2020), and no modulation at 1-3 ms ISIs likely representing intervals too brief for transcallosal propagation (Ni et al. 2020). Short-and long-IHI likely arise from interaction with GABAergic inhibitory interneurons that suppress pyramidal neuron output (Daskalakis et al. 2002; Hui et al. 2020; Ni et al. 2020). At 0 ms ISI, the observed inhibition exceeded M1-elicited long-IHI (50 ms) and was comparable to short-IHI (10 ms), though smaller in magnitude than PMd-elicited inhibition. Since simultaneous stimulation precludes a transcallosal route (Hanajima et al. 2001; Ni et al. 2020), this likely reflects convergence of descending volleys at subcortical sites (e.g., reticular formation, spinal cord; Fisher et al. 2012; Fregosi et al. 2017). The relatively high CS intensities used (∼1 mV; Civardi et al. 2001; Fiori et al. 2017; Ni et al. 2020) may have contributed to this robust inhibition, consistent with prior evidence that IHI magnitude increases with CS intensity (Ni et al. 2009). Future research should systematically vary CS and TS intensities at the 0 ms ISI to clarify the mechanisms underlying non-callosal M1-M1 interactions.

### Limitations

Certain limitations should be acknowledged when interpreting the findings of this study. First, iSP measurements were performed at a single TMS intensity (150% RMT) and contraction level (50% MVC), based on evidence that sufficiently high TMS intensities over M1 and contraction levels reliably elicit iSPs when only the ipsilateral muscle is activated (Chen et al. 2003; Jung and Ziemann 2006; Davidson and Tremblay 2013). However, there is evidence that lower TMS intensities can also elicit comparable iSP magnitudes from M1 (Chen et al. 2003). Future work should investigate whether PMd-and PMv-elicited iSPs demonstrate similar intensity-dependent and contraction-dependent modulation as M1. Second, the iSP assessment from PMd elicited contralateral MEPs, raising the possibility that current spread to ipsilateral M1 partially contributed to the observed inhibition. Yet, these MEPs were significantly smaller than those from direct M1 stimulation, suggesting at least partial premotor origin. This interpretation is further supported by: (1) distinct PMd iSP parameters compared to M1, (2) significant PMv iSPs despite no contralateral MEPs, (3) dual-site IHI using intensities adjusted to avoid contralateral MEPs, minimizing current spread and (4) MNI coordinate estimation showing cortical target separation between PMd, PMv and M1, with distances exceeding the typical current spread of TMS (Opitz et al. 2011). Third, our study would have benefited from examining additional ISIs. Shorter ISIs (4-5 ms) would clarify whether PMd or PMv elicit facilitation similar to M1 at these latencies (Hanajima et al. 2001; Ni et al. 2020). Intermediate ISIs for PMd (between 10 and 50 ms) and longer ISIs for PMv (80 and 150 ms) would provide more complete characterization, particularly given that Fiori et al. (2017) demonstrated PMv facilitation at 80 ms and long-latency IHI at 150 ms. However, we prioritized specific ISIs to compare well-established mechanisms (i.e., short-and long-IHI) alongside novel non-callosal IHI (0 ms ISI) motivated by non-human primate work (Quessy et al. 2016; Côté et al. 2017). Finally, while our non-callosal IHI findings at 0 ms ISI are novel, we can only speculate about convergence sites of subcortical structures (reticular formation, brainstem), the spinal cord, or both. Since transcallosal conduction in humans requires at least 4-5 ms to modulate contralateral M1 (Hanajima et al. 2001; Ni et al. 2020), we are relatively confident the 0 ms ISI inhibition was not transcallosally mediated. Nevertheless, future research should investigate whether simultaneous stimulation from PMd, PMv, and M1 elicits such inhibition via differential or overlapping subcortical and spinal convergence sites.

## Conclusion

This study provides a comprehensive characterization of the differential modulation of contralateral M1 excitability by PMd and PMv in humans. We demonstrate that an iSP can be elicited from PMd and PMv, revealing distinct transcallosal inhibitory connectivity with contralateral M1. Additionally, our dual-site TMS approach mapped short-, long-, and non-callosal interhemispheric inhibitory mechanisms as elicited from PMd, PMv, and M1. Notably, the association between iSP duration and long-IHI magnitude from both PMd and PMv suggests these measures reflect partly overlapping transcallosal mechanisms, bridging our single-pulse and dual-site findings. Our results set the stage for further foundational examination of the mechanisms by which distinct premotor cortical regions, like PMd and PMv, shape M1 output. Finally, our findings of differential connectivity patterns may be important when considering neuromodulation strategies targeting premotor regions in stroke rehabilitation.

## Data availability statement

All main data presented in this manuscript are included within the figures. The datasets produced and analyzed in this study are available from the corresponding author upon reasonable request.

## Author contributions

EA contributed to the methodological development of the study, collected and analyzed the data and wrote the first draft of the manuscript. LC and NH contributed to data collection. AOF contributed to data processing. JLN and ND conceived of the project, contributed to interpretation of data as well as writing and editing the manuscript. All authors edited the manuscript and approved the final version before submission.

