## Supplementary Material for "Dorsal and ventral premotor cortices differentially influence contralateral motor cortex excitability"

| TS | M1 | PMd | PMv |
|---|---|---|---|
| TS Intensity (% MSO) | 55.7 (11.1) | 55.26 (9.9) | 55.5 (10.3) |
| TS Amplitude (mV) | 1.39 (0.39) | 1.35 (0.47) | 1.40 (0.72) |

***Supplementary Table 1: Test Stimulus data***

*Data shown is the peak-to-peak amplitude of the motor evoked potential expressed in mV and intensity of test stimulus expressed in % MSO reported as mean (SD).* *Abbreviations:* ***M1:*** *primary motor cortex,* ***PMd:*** *dorsal premotor cortex,* ***PMv:*** *ventral premotor cortex*
